# Expedited Assessment of Terrestrial Arthropod Diversity by Coupling Malaise Traps with DNA Barcoding

**DOI:** 10.1101/192732

**Authors:** JR deWaard, V Levesque-Beaudin, SL deWaard, NV Ivanova, JTA McKeown, R Miskie, S Naik, K Perez, S Ratnasingham, CN Sobel, JE Sones, C Steinke, AC Telfer, AD Young, MR Young, EV Zakharov, PDN Hebert

## Abstract

1. Monitoring changes in terrestrial arthropod communities over space and time requires a dramatic increase in the speed and accuracy of processing samples that cannot be achieved with morphological approaches.
2. The combination of DNA barcoding and Malaise traps allows expedited, comprehensive inventories of species abundance whose cost will rapidly decline as high-throughput sequencing technologies advance.
3. Aside from detailing protocols from specimen sorting to data release, this paper describes their use in a survey of arthropod diversity in a national park that examined 20,000 specimens representing 2200 species.
4. These protocols can support arthropod monitoring programs at regional, national, and continental scales.

## Introduction

Given unprecedented losses (Lawton & May 1995; Pimm et al. 1995), improved methods to quantify biodiversity are essential, especially for smaller organisms. The melding of two technologies – DNA barcoding and passive, large-scale specimen collection – represents a potential solution. DNA barcoding simplifies and accelerates taxonomic identifications (Hebert et al. 2003; Packer et al. 2009; Cristescu 2014; Joly et al. 2014) by employing the 5.6 million reference sequences in the Barcode of Life Datasystems (BOLD; Ratnasingham & Hebert 2007). Coverage varies for geographic regions and taxonomic groups, ranging from nearly complete for some continental faunas (Hebert et al. 2013; Pentinsaari et al. 2014; Hendrich et al. 2014; Huemer et al. 2014; Rougerie et al. 2014; Zahiri et al. 2017; Blagoev et al. 2015; Gwiazdowski et al. 2015) to sparse for most taxa (Ferri et al. 2009; Hogg et al. 2010; Young et al. 2012; Layton et al. 2014). Because the latter groups include many undescribed species, operational taxonomic units (OTUs) must be employed to quantify their diversity. DNA barcoding represents a dramatic advance for such analysis because the Barcode Index Number (BIN) system (Ratnasingham & Hebert 2013) provides an objective approach for OTU delineation that is coupled to a persistent registry. Since BINs are strong proxies for Linnaean species (Hausmann et al. 2013; Ratnasingham & Hebert 2013; Zahiri et al. 2014; Blagoev et al. 2015), biodiversity assessments based on BINs can be implemented for groups that lack well-developed taxonomy.

The Malaise trap (Malaise 1937) has gained popularity for assessing terrestrial arthropod communities (Karlsson et al. 2005) because it collects large samples with little effort. However, the subsequent identification is a substantial challenge as a week-long collection often includes more than 1000 specimens representing several hundred species. Moreover, because many species are only represented by a few specimens, it is important to identify every individual. Conversely, very common species can consume considerable effort, particularly if they belong to a closely allied group of taxa whose members are difficult to discriminate morphologically. DNA barcoding breaks this taxonomic barrier as it can rapidly identify all individuals.

While the analysis of bulk samples through DNA metabarcoding (Hajibabaei et al. 2011; Taberlet et al. 2012; Yu et al. 2012; Ji et al. 2013; Gibson et al. 2014; Leray & Knowlton 2015) greatly reduces analytical costs, it has two limitations. It cannot maintain the link between each specimen and its COI sequence, which inhibits extending the DNA barcode reference library, and cannot determine species abundances.

This study describes a protocol for rapid biodiversity assessments which employs DNA barcoding and passive specimen trapping. Its effectiveness is demonstrated by describing a survey that examined 20,000 specimens representing over 2200 species from Point Pelee National Park.

## Materials and Methods

### Specimen Collection and Processing

A Townes-style Malaise trap was deployed in a cedar-savannah habitat at Point Pelee National Park in southwestern Ontario from May 2 until September 19, 2012. Each sample was collected in a 500 mL plastic Nalgene bottle that was filled with 375 mL of 95% ethanol and then attached to the trap head (Fig. 1 F2). The catch was harvested weekly and placed in 500 mL of fresh ethanol before storage at -20°C until it was analyzed at the Centre for Biodiversity Genomics (CBG; http://www.biodiversitygenomics.net).

**Figure 1.**
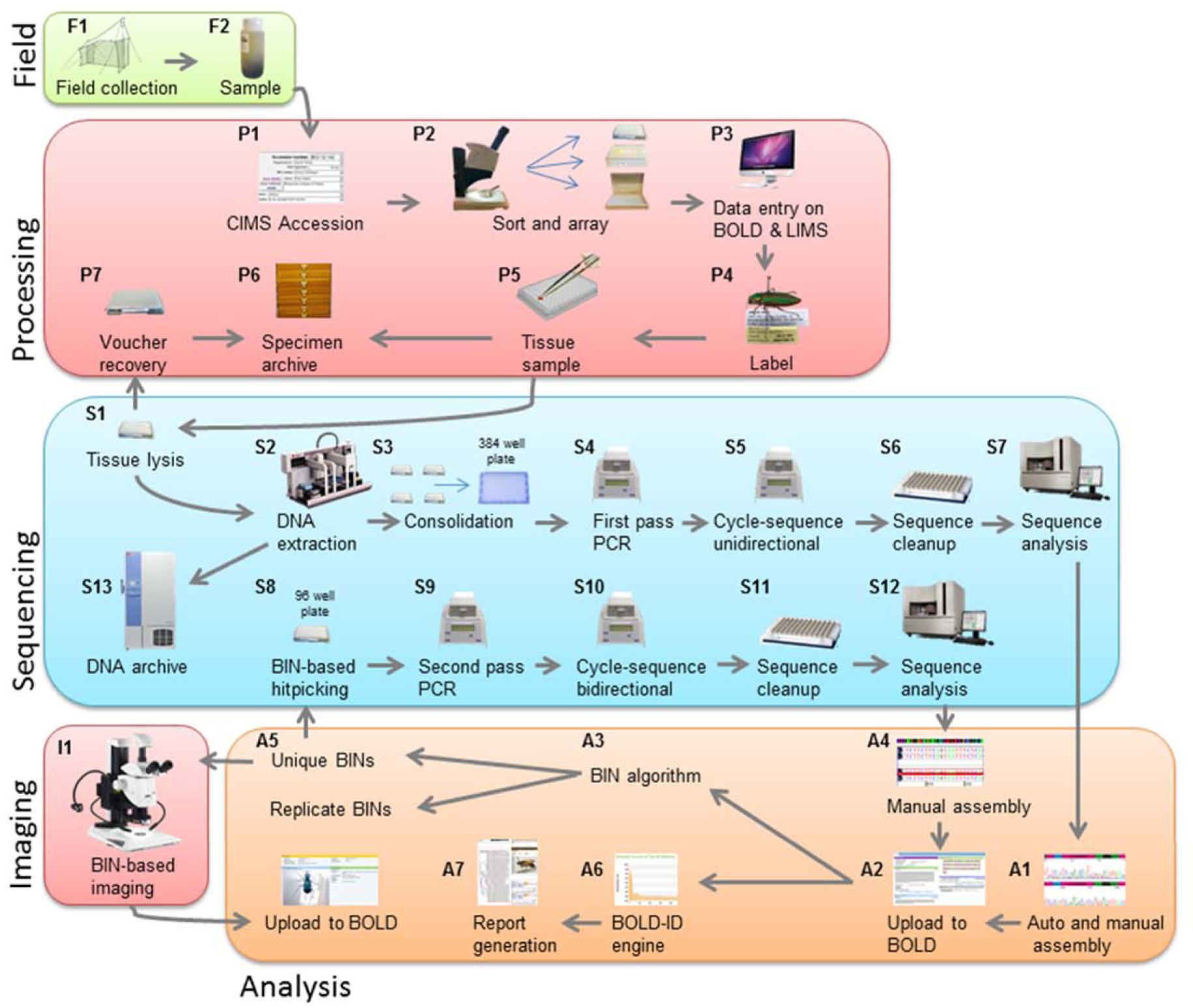
Workflow for biodiversity monitoring through DNA barcoding.

Each weekly sample was accessioned and its collection data entered into an Access-based Collection Information Management System (CIMS; Fig. 1 P1). Samples collected in odd-numbered weeks (1, 3, 5…) were sequenced while the others were archived. The first stage in the analysis involved decanting excess ethanol, and pouring the specimens into a sorting dish. Specimens were then partitioned by size (small, medium and large) and assigned to a taxonomic order. Large specimens (>5mm) were pinned while medium and small were retained in ethanol. Three samples contained >300 specimens of a particular OTU. In these cases, 24 specimens were barcoded while the others were counted and recorded. After sorting, specimens were arrayed in batches of 95 plus one control (Fig. 1 P2), mirroring the 8 x 12 format of 96 well microplates. Specimens of different orders were only combined when necessary to fill a plate (Table 1). Large specimens were placed in Schmitt boxes with an 8 x 12 grid marked on their foam base, while medium specimens were placed in Matrix storage tubes (Thermo Scientific; Fig. 1 P2), and small specimens were placed directly in 96-well microplates (Eppendorf; Fig. 1 P2). Each container was given a unique identifier (Root Plate ID) and likewise, each specimen within the container was given a unique identifier reflecting its position in it (Sample ID). The unique identifiers and collection data for each specimen were uploaded to BOLD (Ratnasingham & Hebert 2007; Fig. 1 P3) with records for each sample placed in a separate project to allow easier comparison among weeks. Once this was completed and BOLD Process IDs were generated, labels were printed and affixed to large and medium specimens while small specimens did not require individual labels (Fig. 1 P4). A small tissue sample was then removed from each large and medium specimen and placed into a microplate destined for DNA extraction (Fig. 1 P5). Small specimens did not require tissue sampling as they were already in microplates. Each microplate was then submitted for sequence analysis and its progress through the analytical chain was tracked with a Laboratory Information Management System (BOLD-LIMS).

**Table 1.**
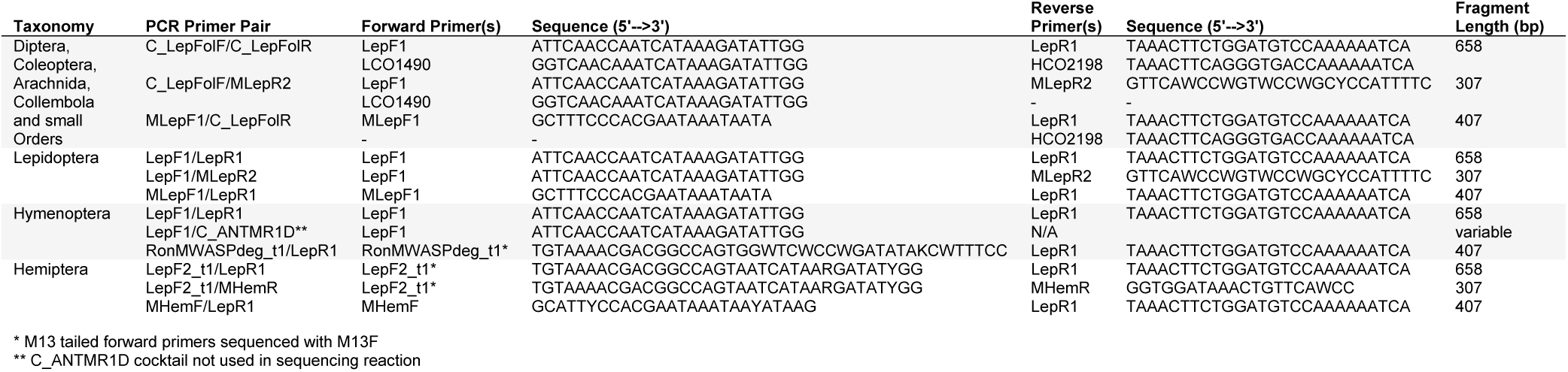
Three first pass PCR primer pairs used to amplify the 658 bp barcode region of COI and eight second pass PCR primer pairs used to amplify two smaller fragments (307 bp, 407 bp or variable) of this gene region. Arachnida, Coleoptera, Collembola, Diptera and small orders share the same first and second pass primer pairs whereas Lepidoptera and Hymenoptera only share the first pass primer pair as they were processed with second pass primers specific to each order. Hemiptera were processed with first and second pass primers unique to this order. The listed primer was used for sequencing unless indicated by an asterisk.

### DNA Barcode Analysis

Sequence analyses were conducted at the Canadian Centre for DNA Barcoding (CCDB; http://www.ccdb.ca). An automated, silica membrane-based DNA extraction protocol (Ivanova et al. 2006) was performed in 96-well microplate format using a 3 μm glass fibre over 0.2 μm Bio-Inert membrane filter plate (Pall Corporation). To maximize DNA yield, tissue lysis was performed overnight at 56°C before DNA extraction (Fig. 1 S1 and S2). Subsequent PCR amplification of the COI barcode region was performed in 384-well plate format as this allowed a 50% reduction in reagent volumes from earlier methods (Hajibabaei et al. 2005; deWaard et al. 2008, Wilson 2012). This protocol involved consolidating aliquots of DNA extracts from four 96-well microplates into a 384-well PCR plate containing PCR master mix using a Biomek FX workstation (Beckman-Coulter; Fig. 1 S3) and ensured arthropod orders were processed with the same primer pair. The total PCR reaction volume was 6 μL: 3 μL of 10% D-(+)-trehalose dihydrate for microbiology (≥99.0%; Fluka Analytical), 0.92 μL of ultra-pure water (Hyclone, Thermo Scientific), 0.60 μL of 10× PlatinumTaq buffer (Invitrogen), 0.30 μL of 50 mM MgCl_2_ (Invitrogen), 0.06 μL of each primer, 0.03 μL of 10 mM dNTP (KAPA Biosystems), 0.03 μL of 5 U/μL PlatinumTaq DNA Polymerase (Invitrogen), and 1 μL of DNA template. Table 1 details the primer pairs used on the first pass. All PCR reactions employed the same thermocycling parameters: 94°C for 1 min, 5 cycles at 94°C for 40 sec, 45°C for 40 sec, 72°C for 1 min, followed by 35 cycles at 94°C for 40 sec, 51°C for 40 sec, 72°C for 1 min, and a final extension at 72°C for 5 min (Fig. 1 S4).

PCR products were diluted 1:4 with molecular grade water and then unidirectionally sequenced using the appropriate reverse primer (Table 1). Sequencing was also completed in 384-well format (Fig. 1 S5) to reduce costs. The total sequencing reaction volume was 5.5 μL: 0.14 μL of BigDye terminator v3.1 (Applied Biosystems), 1.04 μL of 5X sequencing buffer [400 mM Tris-HCl pH 9.0 + 10 mM MgCl_2_ (Invitrogen)], 2.78 μL of 10% D-(+)-trehalose dihydrate from *Saccharomyces cerevisiae* (≥99%; Sigma-Aldrich), 0.48 μL of ultra-pure water (Hyclone, Thermo Scientific), 0.56 μL of primer; and 0.5 μL of diluted PCR template was added with a Biomek FX robot. All sequencing reactions employed the same thermocycling protocol: 96°C for 1 min followed by 15 cycles at 96°C for 10 sec, 55°C for 5 sec, 60°C for 1.25 min, followed by 5 cycles at 96°C for 10 sec, 55°C for 5 sec, 60°C for 1.75 min, then 60°C for 15 sec followed by 15 cycles at 96°C for 10 sec, 55°C for 5 sec, 60°C for 2 min and a final extension at 60°C for 1 min (Fig. 1 S6). An automated, magnetic bead-based sequencing cleanup method was employed in 384-well microplates using PureSEQ (ALINE Biosciences) on a separate Biomek FX robot before sequencing on an ABI 3730xL DNA Analyzer (Applied Biosystems; Fig. 1 S7).

Trace files were manually uploaded to BOLD and were automatically assessed for quality based on predefined parameters (Ratnasingham & Hebert 2007). Trace files that received medium and high-quality assessments were automatically trimmed and edited by the BOLD platform. Those deemed to be low quality or classified as failed reads were ignored. Trimming was performed using a sliding window approach, discarding leading and trailing segments of the sequence that had more than 4 bp with a QV lower than 20 in a window of 20 bp. All sequences with less than 500 bp in the barcode region (the threshold for BIN assignment; see below) were manually edited with CodonCode v. 3.0.1 (CodonCode Corporation) to see if additional sequence information could be recovered (Fig. 1 A1).

The initial PCR failed to generate an amplicon from some DNA extracts, likely reflecting DNA degradation or low primer affinity. These failures were hitpicked to assemble new destination 96-well microplates of DNA extracts (Fig. 1 S8), which were subjected to another round of PCR employing primers that generated two overlapping COI (307 bp, 407 bp) amplicons (Table 1; Fig. 1 S9). A Biomek NX Span 8 workstation (Beckman-Coulter) was used to hitpick the failed samples into new plates. This ‘failure tracking’ was supported by data generated by the BOLD-LIMS. The original DNA plates were scanned to identify all specimens that failed to generate a barcode compliant sequence. The well coordinates of these failed samples in the source and destination microplates were generated for input into a Biomek NX robot. The newly configured microplates were then processed through two PCR reactions followed by bidirectional sequencing and manual assembly as part of the failure tracking protocol (Fig. 1 S10, S11, S12 and A4). Failure-tracking PCR reactions were carried out in 96-well microplates. The total PCR reaction volume was 12.5 μL: 6.25 μL of 10% D-(+)-trehalose dihydrate for microbiology (≥99.0%; Fluka Analytical), 0.125 μL of ultra-pure water (Hyclone, Thermo Scientific), 2.5 μL of 5× KAPA Taq HotStart Buffer (KAPA Biosystems), 1.25 μL of 25 mM MgCl_2_ (Invitrogen), 0.125 μL of each primer, 0.0625 μL of 10 mM dNTP (KAPA Biosystems), 0.0625 μL of 5 U/μL KAPA Taq HotStart DNA Polymerase (KAPA Biosystems), and 2 μL of DNA template. Failure-tracking sequencing reactions were also carried out in 96-well microplates. PCR products were diluted 1:5 and bi-directionally sequenced. The total sequencing reaction volume was 11 μL: 0.25 μL of BigDye terminator v3.1 (Applied Biosystems), 1.875 μL of 5X sequencing buffer [400 mM Tris-HCl pH 9.0 + 10 mM MgCl_2_ (Invitrogen)], 5 μL of 10% D-(+)-trehalose dihydrate from *Saccharomyces cerevisiae* (≥99%; Sigma-Aldrich), 0.875 μL of ultra-pure water (Hyclone, Thermo Scientific), 1 μL of primer; and 2 μL of diluted PCR template.

The final step in barcode analysis involved a second round of ‘BIN hitpicking’ to ensure that each BIN was represented, whenever possible, by five individuals with bidirectional sequence coverage. BIN information on BOLD was utilized in conjunction with the BOLD-LIMS to select representatives of each BIN with <5 individuals with bidirectional coverage (Fig. 1 A5) and instructions were automatically generated for the Biomek NX Span 8 workstation. The hitpicked destination DNA microplates were then processed through the PCR to bidirectional sequencing steps (Fig. 1 S8 to S12), manually edited (Fig. 1 A4) and uploaded to BOLD (Fig. 1 A2).

### Data Release and Barcode Index Numbers

Specimen and sequence data are available on BOLD (Fig. 1 A2) in the public dataset DS-PPNP12 entitled " Point Pelee National Park Malaise Trap Program 2012" (http://dx.doi.org/10.5883/DS-PPNP12). The record for each specimen includes its date and locality of collection, its taxonomic assignment, and voucher specimen details. If its barcode was recovered, the specimen record also includes trace files, quality scores, its sequence, and corresponding GenBank accession. After final validation, the specimen data were also uploaded to the Global Biodiversity Information Facility (GBIF) as a Darwin Core Archive (Wieczorek et al. 2012) via the Integrated Publishing Toolkit (Robertson et al. 2014) on the Canadensys portal and are available at http://dx.doi.org/10.5886/qzxxd2pa. A condensed version of the data is available in Appendix S1.

The source specimen for each sequence that met quality checks was designated a BIN by the Refined Single Linkage (RESL) algorithm on BOLD (Ratnasingham & Hebert 2013; Fig. 1 A3). The requirements for BIN membership are >=500 bp coverage of the barcode region between positions 70 and 700 of the BOLD alignment, <1% ambiguous bases, and the absence of a stop codon or contamination flag. Alternatively, specimens can gain BIN assignment without formal membership if the sequence is 200–500 bp and unambiguously matches an existing BIN member. RESL runs monthly on all qualifying barcode sequences in BOLD which currently totals 5.74 million specimens and 0.52 million BINs (September 2017). The BIN designations generated through this approach are transparent, reproducible, and globally accessible through DOI-designated ‘BIN pages’ that collate the specimen and sequence information of its members (e.g., *Danaus plexippus* http://dx.doi.org/10.5883/BOLD:AAA9566).

### Archiving and Imaging

All voucher specimens are archived in the collection (BIOUG) at the CBG where they are available for taxonomic study (Fig. 1 P6). Large pinned specimens were assigned to an archive location using BIOUG’s CIMS and transferred to a drawer in the dry collection. Each medium-sized specimen was retained in its storage tube in the Matrix box, assigned an archive location, and stored in BIOUG’s fluid collection. Small specimens were returned from the CCDB after voucher recovery (Porco et al. 2010; Fig. 1 P7), retained in their microplates, and archived in BIOUG’s fluid collection. All residual DNA extracts are stored in the DNA Archive at the CBG (Fig. 1 S13), where they are available for further sequence characterization.

Once sequence analysis was complete and specimens were designated BINs, up to three representatives of each BIN were photographed (Fig. 1 I1) by employing a database query to recognize BINs lacking an image. Specimens were photographed at high resolution and the images were made accessible through both specimen and BIN pages under Creative Commons (BY-NC-CA) license.

### Taxonomic Assignment and Data Analysis

Following BIN designation, every record received a taxonomic assignment based upon querying BOLD (Fig. 1 A6). If the record’s BIN contained specimens identified to a single family, genus or species by a taxonomic expert, it received this identification. Records assigned to a new BIN were queried through the BOLD Identification Engine (http://www.boldsystems.org/index.php/IDS_OpenIdEngine). If the result was a close match (<10% divergence for family, <5% for genus) and the query sequence fell within a monophyletic cluster of BINs assigned to a particular genus or family, the record was assigned to this taxon and confirmed morphologically. All assignments were further validated using the taxon ID tree (Fig. S1) along with matching specimen images (Fig. S2). Any anomalies in tree topology were investigated by retrieving the vouchered specimen and ensuring that all ancillary data on BOLD were correct (including the specimen image and preliminary identification). If the sequence was revealed as representing a contamination event, it was flagged, tagged on BOLD as a contamination, and removed from the analysis and its BIN page.

The final stage of the workflow involved report generation (Fig. 1 A7) aided by the varied functions on BOLD for calculating summary statistics. As well, supplementary analyses were performed to demonstrate the utility of the protocol for rapid biodiversity assessment. To explore the completeness of the inventory, sample- and individual-based accumulation curves were computed using EstimateS 9.1.0 software (Colwell 2013). In addition to constructing a curve for observed species, one was also calculated using the nonparametric species richness estimator Chao 2 (Chao 1987). For all analyses, curves were computed as the mean of 1000 randomized species accumulation curves without replacement. As another measure of completeness, log-normal abundance plots were calculated using the software product R, version 3.1.1 (R Development Core Team) and the vegan package (Oksanen et al. 2013). R was also used to explore and visualize the taxonomic composition of the collection, the distribution of specimens and BINs over the season, their patterns of relative abundance, the incidence of unique and rare BINs, and the turnover of BINs among samples and across time.

## Results

### DNA Barcode Analysis

All specimens in the ten samples were processed except for three abundant OTUs, each from a different sample (week 3: 8,595 specimens of a chironomid; week 5: 313 specimens of a chironomid; week 9: 334 specimens of a sarcoptiform mite). After their exclusion, the number of specimens per sample averaged 1,907 (range = 814–3,795) (Fig. 4). In total, 21,194 specimens were processed from the ten samples with first pass analysis generating sequences from 81.6% of them (17,300). The second pass analysis recovered another 1885 sequences, bringing the success rate to 90.5% (19,185). Aside from these records, 144 sequences were found to be contaminants and another eight possessed stop codons (Fig. 2). Barcode recovery varied among taxa with Acari displaying the lowest success with just 48.0% of specimens generating a barcode compliant sequence. There was also evidence of a taxonomic bias in the 309 (1.6%) specimens that were either destroyed or unrecoverable after analysis with most being small, soft-bodied Hemiptera (33.7%), Diptera (24.3%) or Acari (21.7%).

**Figure 2.**
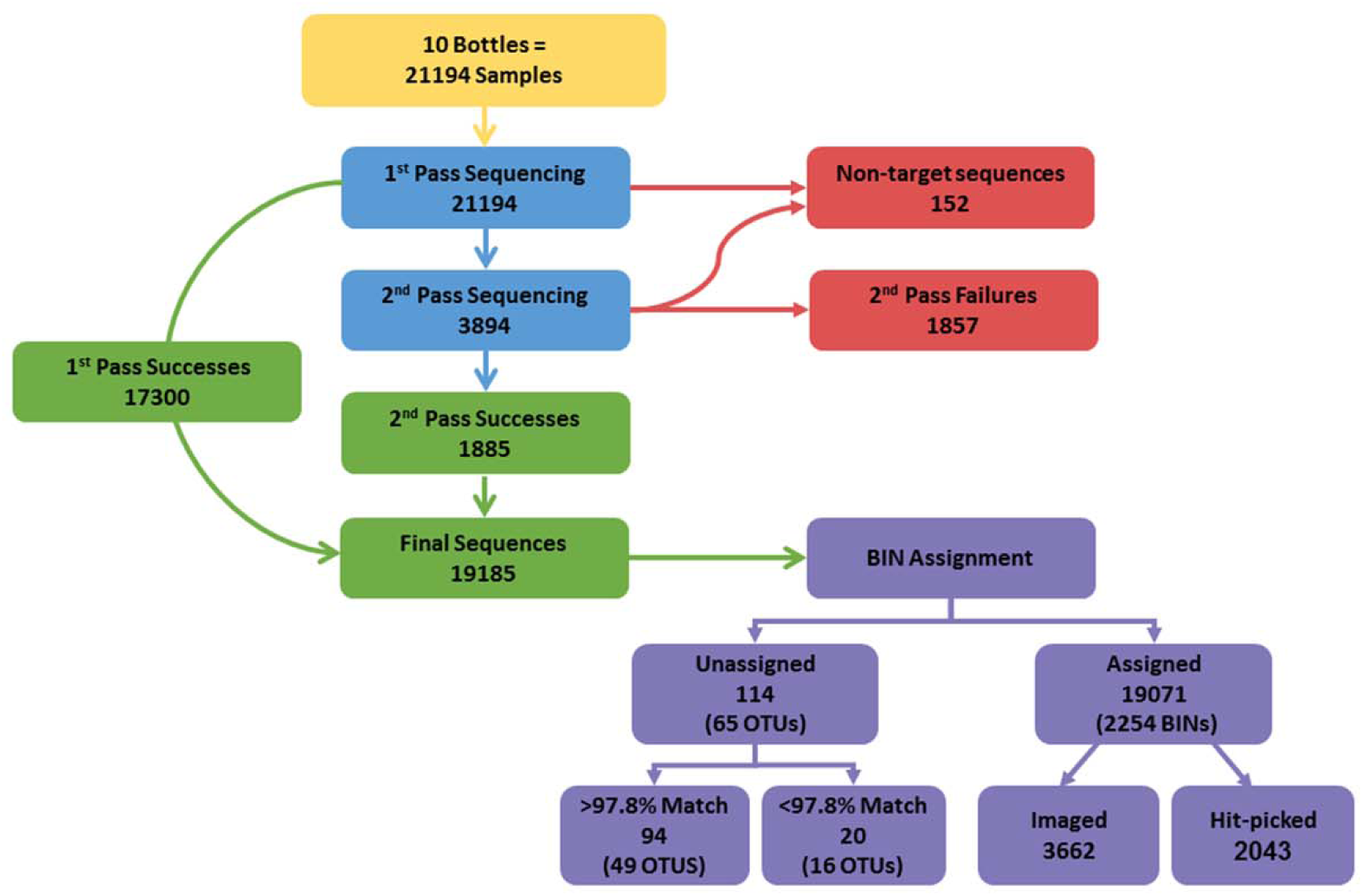
Flowchart showing the success in sequence recovery from 21,194 specimens of arthropods in ten Malaise trap samples.

**Figure 3.**
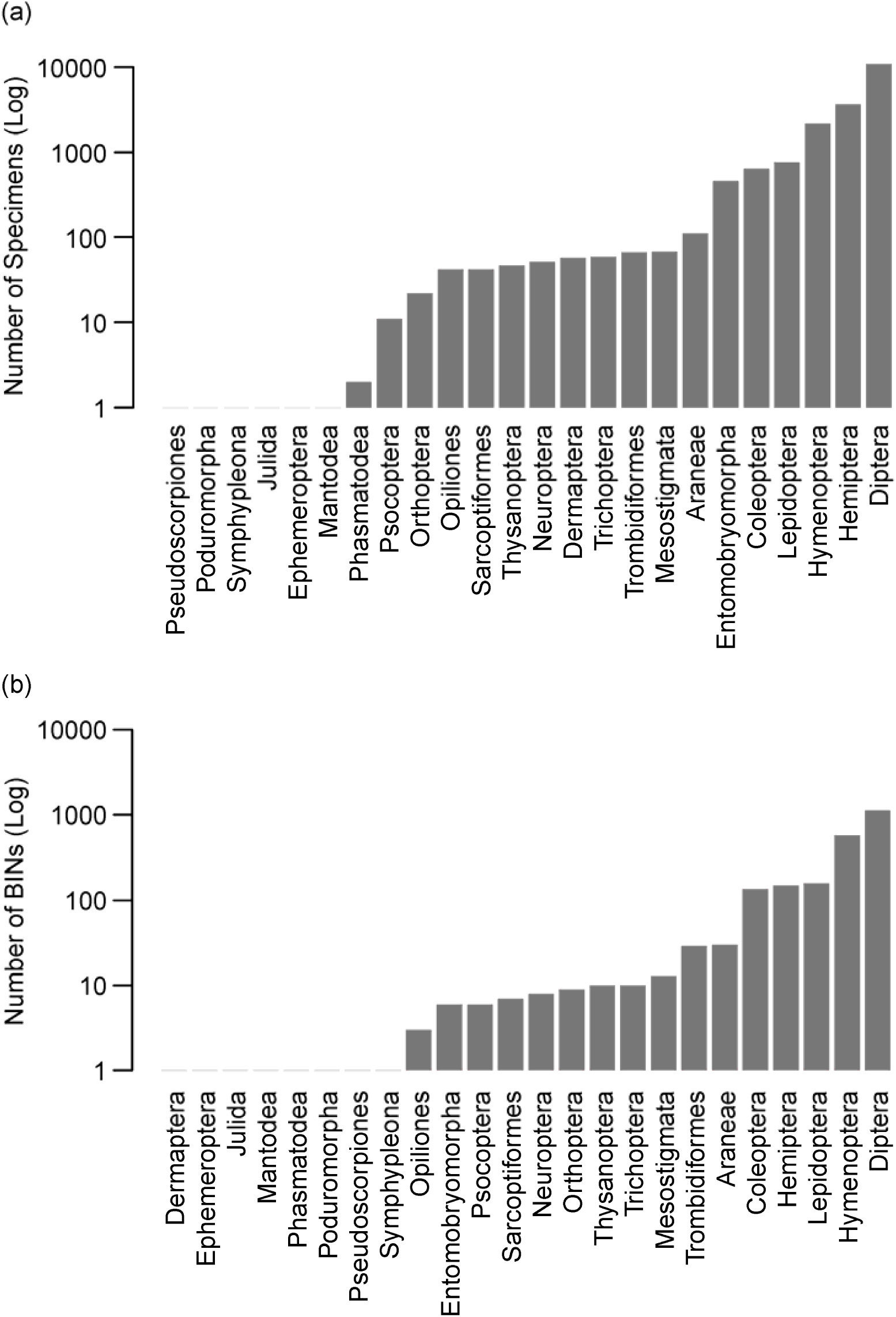
Taxonomic breakdown of the Malaise trap samples by (a) specimens and (b) BINs.

**Figure 4.**
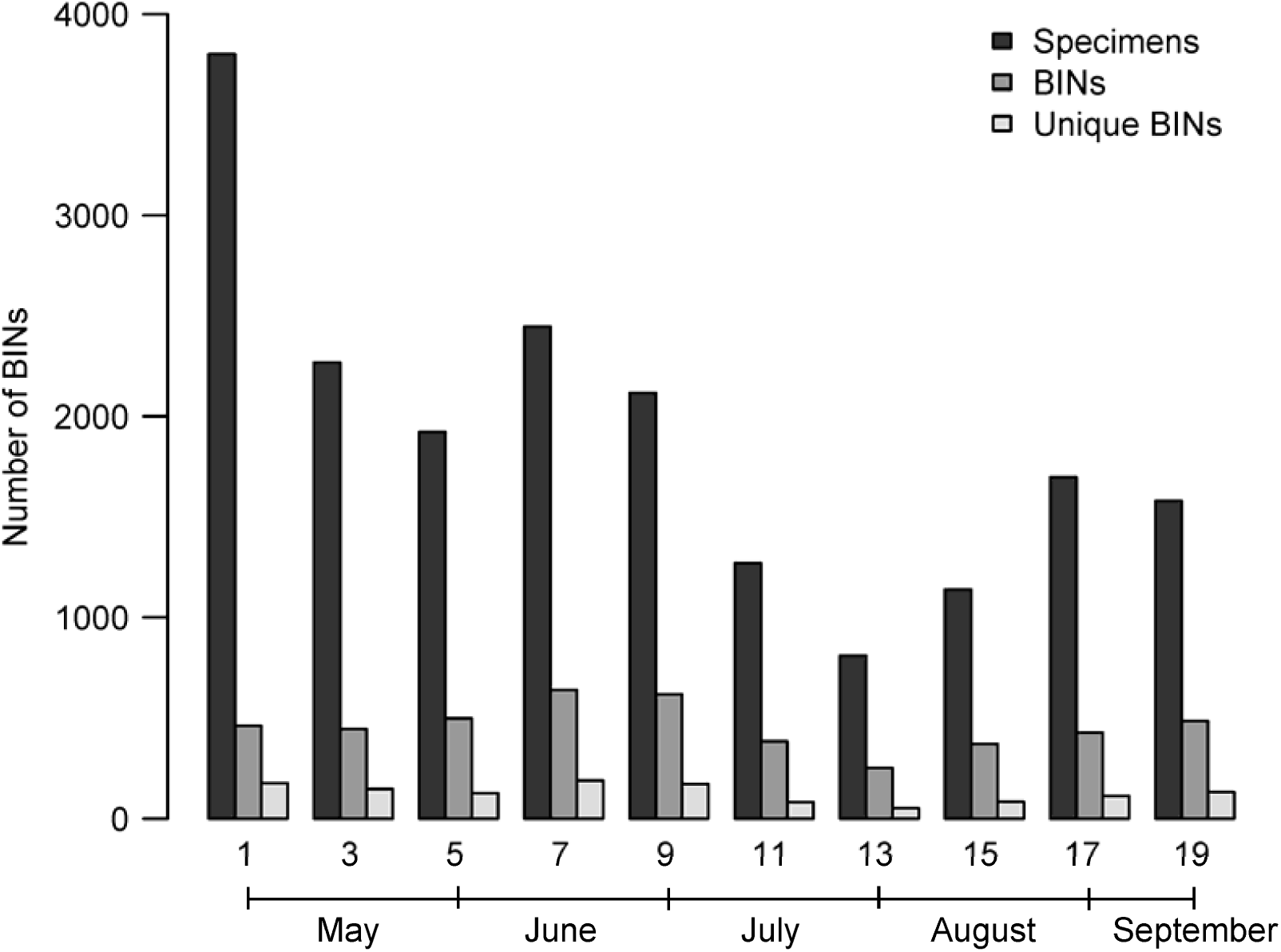
The number of specimens and BINs in ten Malaise trap samples from Point Pelee National Park.

### Specimen and BIN Analyses

Among the specimens that generated a sequence, most (99.4%) received a BIN designation (n = 19,071) (Fig. 2). From these, representatives of 2,403 new BINs were ‘BIN hitpicked’ to acquire a bidirectional sequence for the barcode reference library on BOLD and 3,662 BINs were imaged (mean = 1.6 images/BIN). The 114 sequences that failed to meet the criteria for BIN designation were run through the stand-alone version of the RESL algorithm (using the function ‘Cluster sequences’ on BOLD) to estimate the number of additional OTUs (or species) represented. This analysis revealed 65 OTUs; 49 were likely matches to known BINs (>97.8% match) while 16 were probably new (< 97.8% similarity).

All subsequent analyses considered the 19,071 specimens with a BIN designation. They included taxa belonging to four classes and 25 orders (Fig. 3). Diptera were dominant comprising 57.0% of the specimens (Fig. 3a) and 49.7% of the BINs (Fig. 3b). Hymenoptera was also very diverse with the third highest percentage of specimens (11.3%) and the second highest BIN count (25.3%).

In total, 2,254 BINs were present in the ten samples with an average of 458 per sample (range = 253–640) (Fig. 4). Most were uncommon; 47.6% (1,074) of the BINs were represented by a single specimen while only 36 (1.6%) had >100 specimens (Fig. 5). There was positive correlation between the number of individuals in a sample and the number of BINs unique to it (R^2^ = 0.69, p = 0.003, Fig. 6), reinforcing the prevalence of rare BINs and the effort required to discover them.

**Figure 5.**
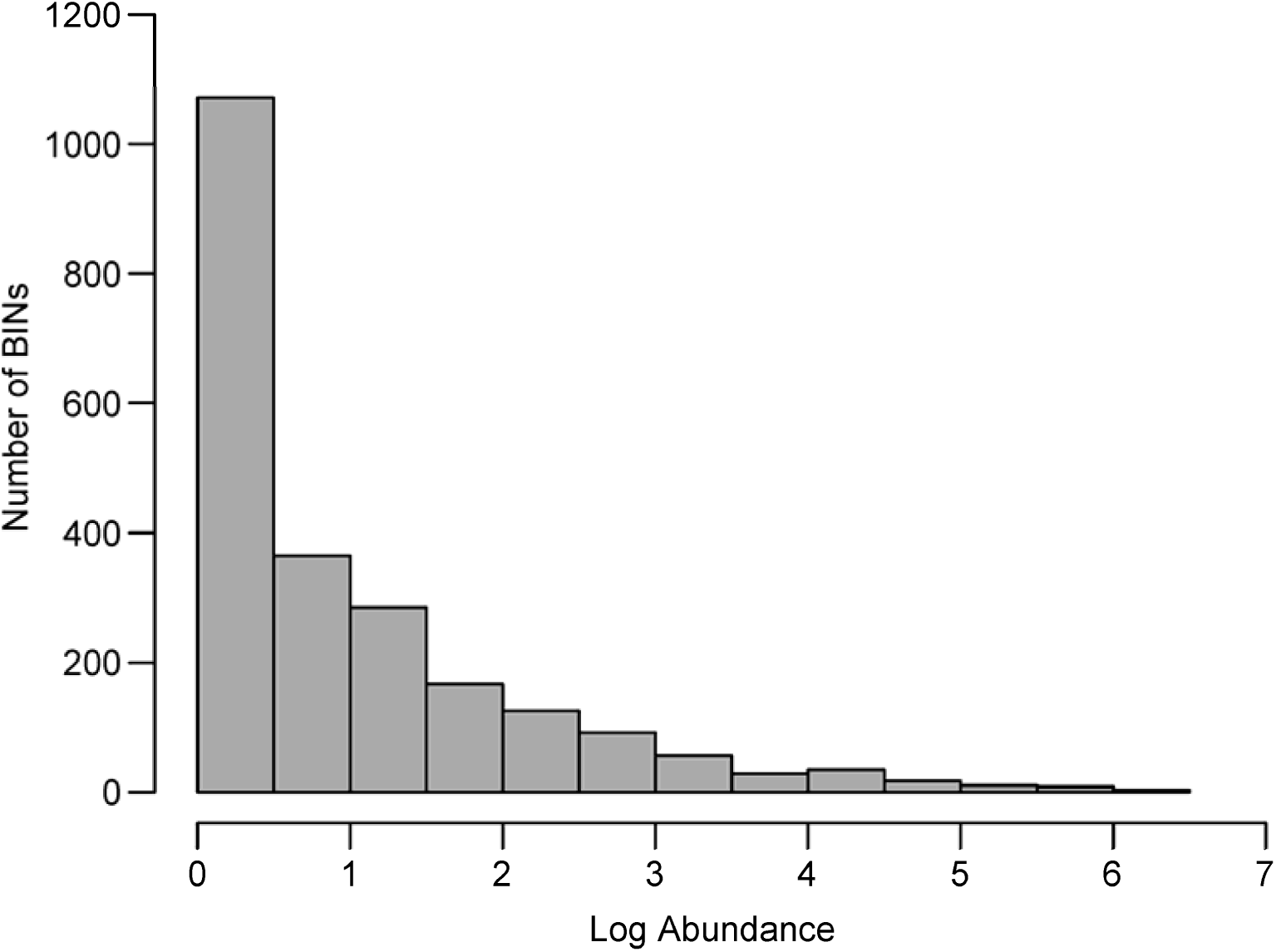
Relative species abundance plot for the ten Malaise trap samples.

**Figure 6.**
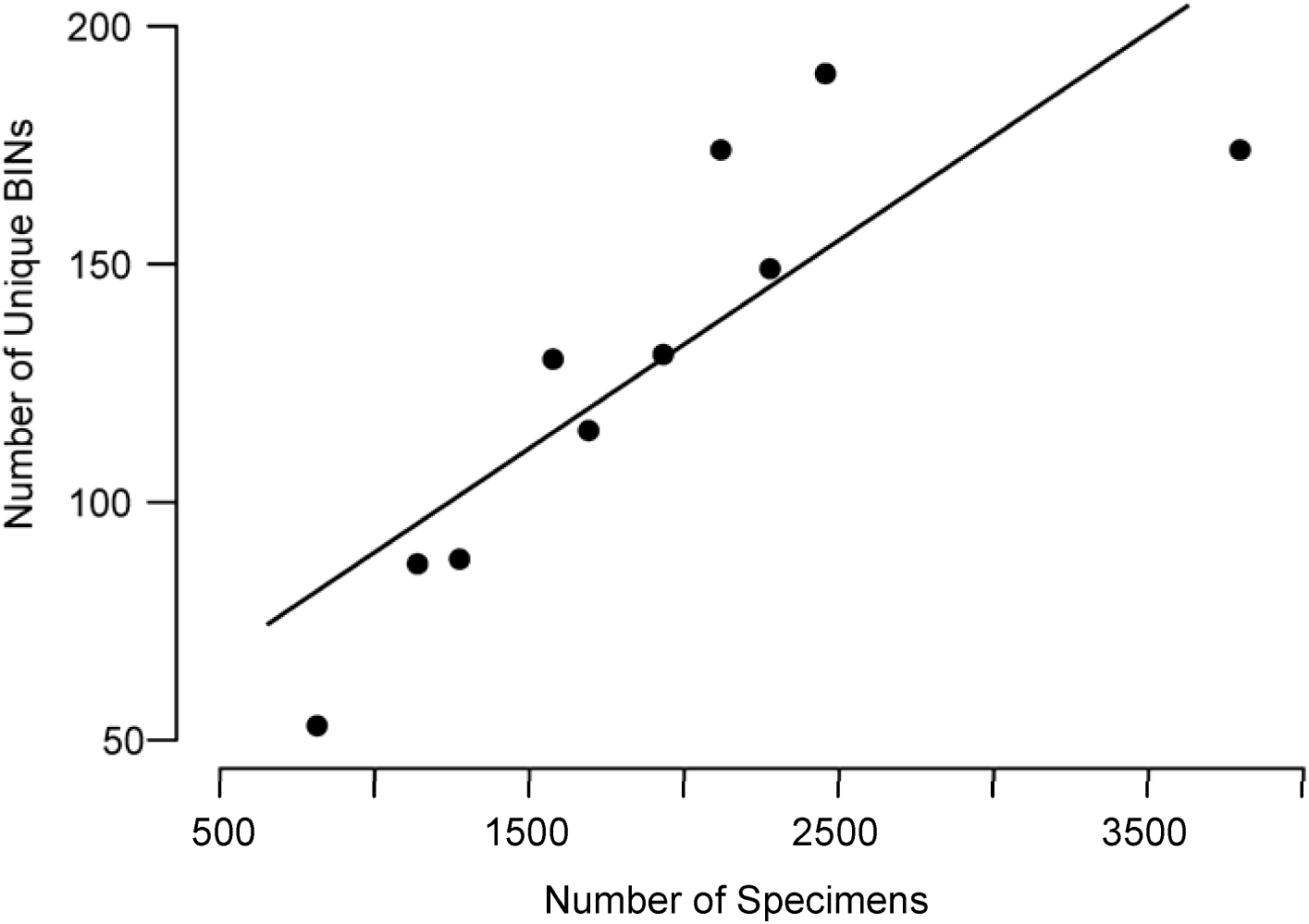
Relationship between the number of specimens in each of ten samples from Point Pelee National Park and the number of BINs unique to it.

### Species Richness and Turnover

Species richness extrapolation based on the (Preston) log-normal species distribution indicated that complete sampling of the Malaise-trappable arthropod fauna at Point Pelee would reveal about 5,700 BINs, roughly double the observed number (Fig. 7). BIN accumulation curves based on Chao 1 suggested a slightly lower count with an estimate of 3,866 (± 133) BINs based on specimens (Fig. 8a) and 4,070 (± 138) based on samples (Fig. 8b). These three estimators suggest the site inventory is roughly 39.6–58.5% complete.

**Figure 7.**
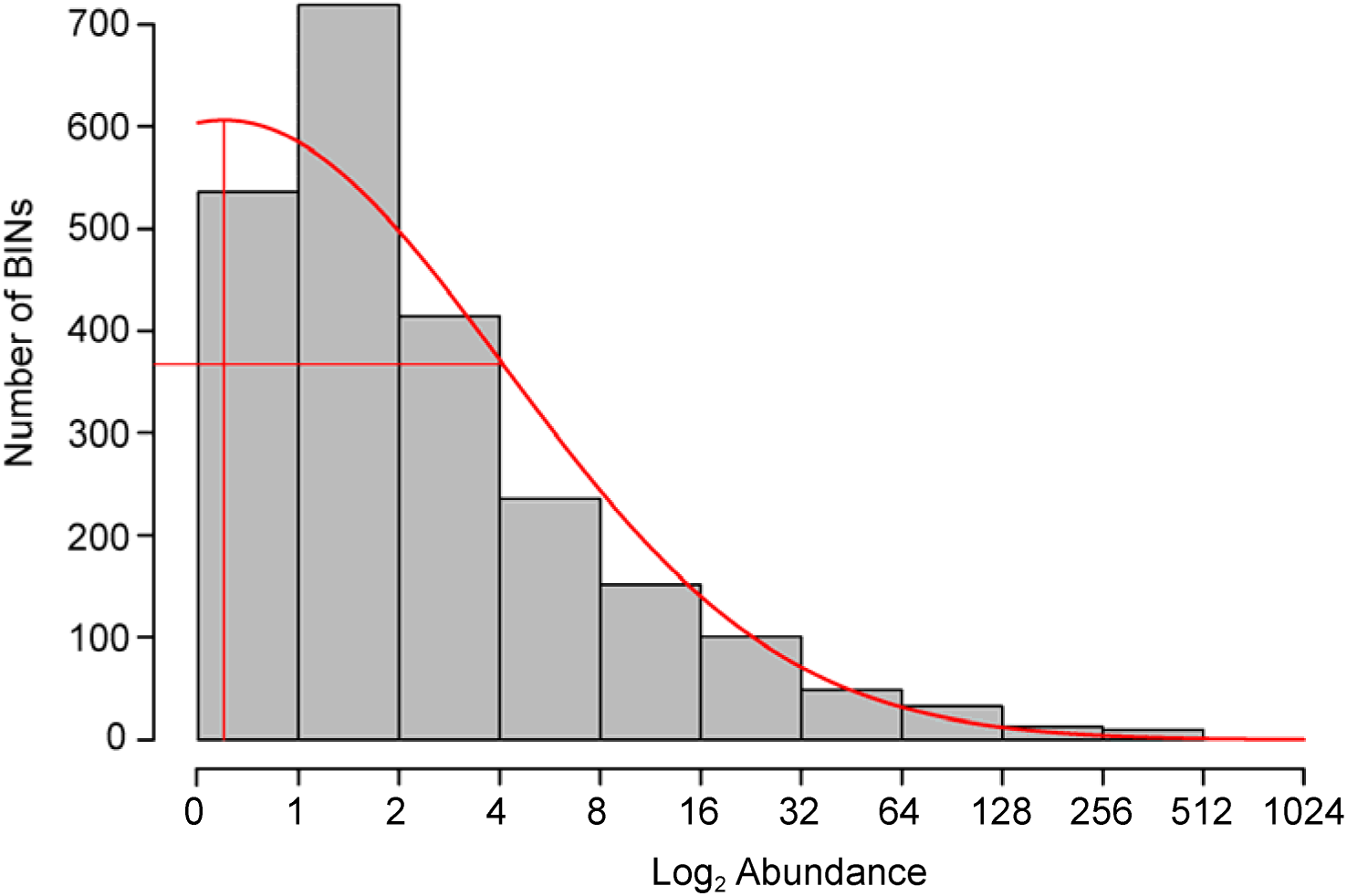
Preston plot with veil line and extrapolation based upon the abundance data for the taxa represented among the 19,071 arthropods that generated a sequence.

**Figure 8.**
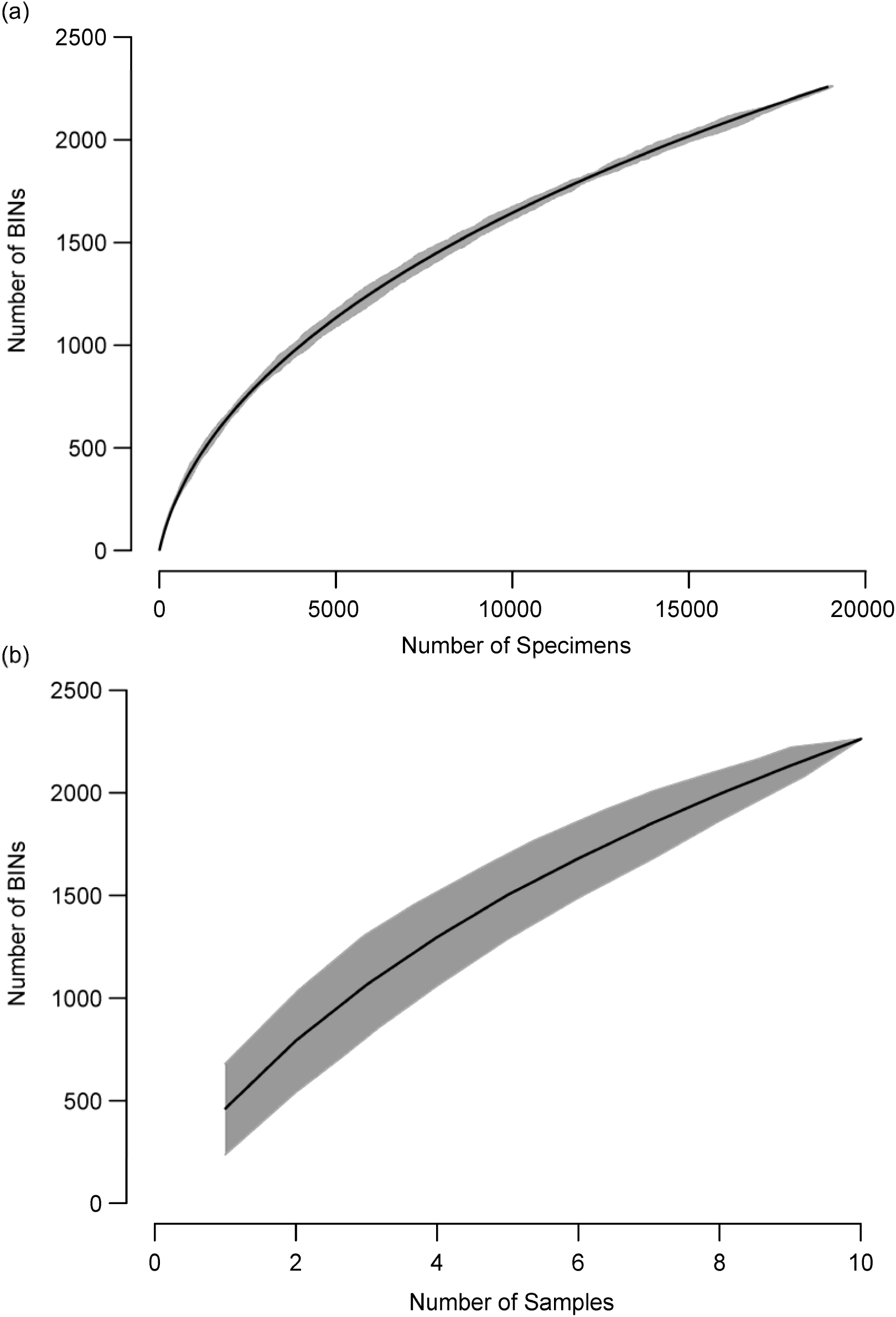
BIN accumulation curves for the 19,071 arthropods from Point Pelee National Park estimated with (a) specimens and (b) samples.

Individual samples contained an average of 458 BINs, but their similarity was low (mean shared BINs = 0.33; mean Jaccard index = 0.16). The proportion of shared BINs (for adjacent and non-adjacent samples) increased as the season progressed (Fig. 9a) and decreased with the interval between samples (R^2^ = 0.52, p << 0.001, Fig. 9b) with similarity values (Jaccard index) halved in 81.1 days. For example, only 99 BINs were shared between weeks 1 and 19, samples that contained 461 and 486 BINs, respectively. By comparison, samples from weeks 7 and 9 (641 and 619 respectively) shared 266 BINs.

**Figure 9.**
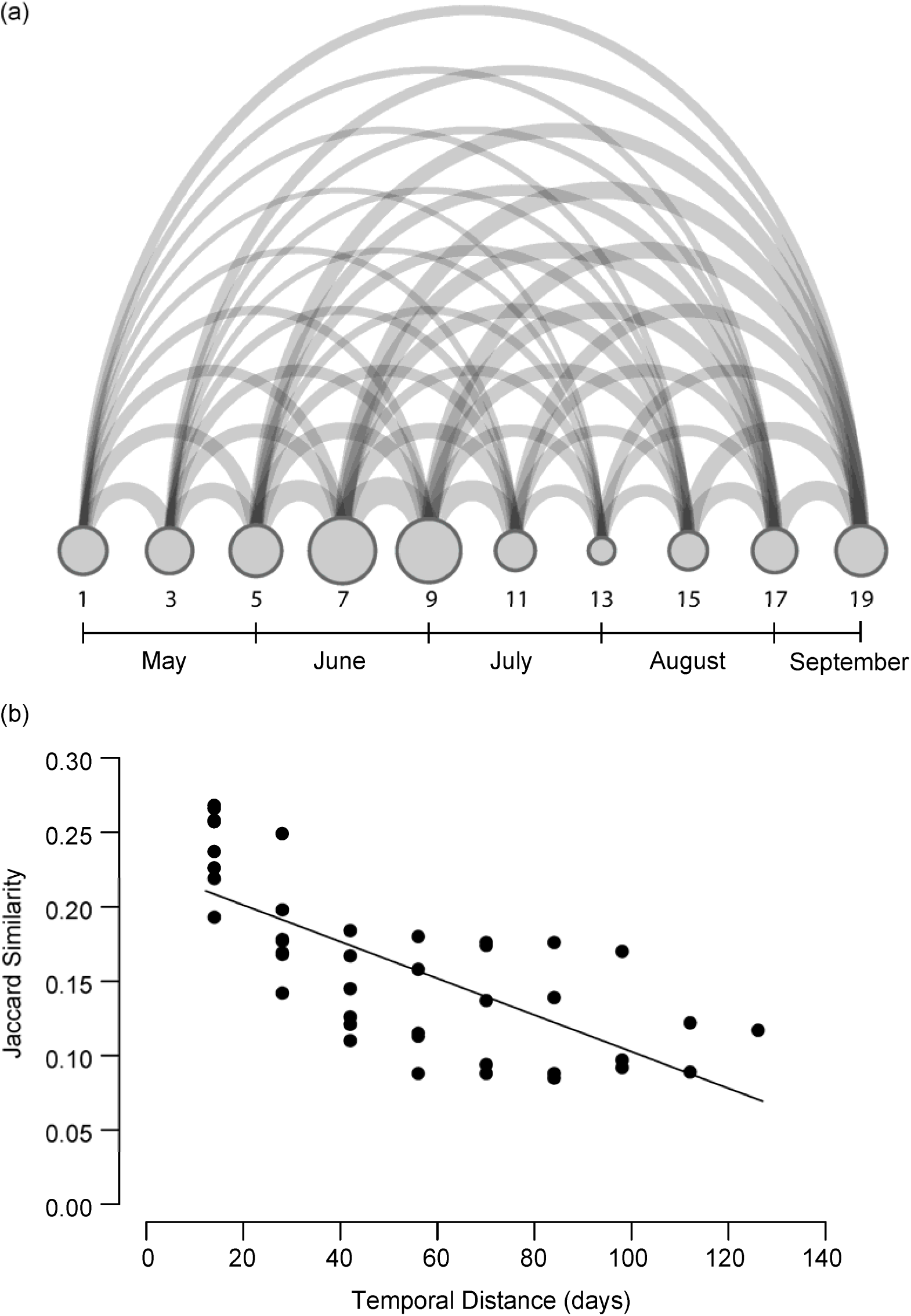
Species overlap between the ten Malaise trap samples, shown (a) in chronological order with the size of each node proportional to the number of BINs in a sample while the width of each arcs reflects BIN overlap between samples. (b) as a comparison of BIN overlap with time between samples.

### Taxon Diversity and Abundance

The Cecidomyiidae (351), Ichneumonidae (127) and Chironomidae (113) included the most BINs while the Chironomidae (10,827), Cicadellidae (3,070), and Cecidomyiidae (1,919) were represented by the most specimens. The most abundant BINs were BOLD:AAG2868 (Cicadellidae: *Empoasca fabae*), BOLD:AAB7030 (Chironomidae: *Chironomus* sp.) and BOLD:AAV0161 (Cicadellidae: *Erythroneura bakeri*) with 555, 446 and 431 specimens respectively. Each of these species and many of the abundant taxa had closely-related allies, often morphologically indistinguishable and in low frequency, making oversampling unavoidable without risking the oversight of some species.

### New and Existing Inventories

Three quarters of the specimens (n = 14,313/19,071) with a sequence gained a genus- or species-level taxonomic assignment following their comparison with records on BOLD. They represented 58.6% of all BINs (n = 1320); the other BINs were assigned to a subfamily or family. A few species were represented by more than one BIN [e.g., Araneae: Thomisidae: *Xysticus pellax* represented by BINs BOLD:ACE4932 and BOLD:ACE4935], but most species (95.5%) showed perfect correspondence between a single taxon name and a single BIN.

By comparison, a 40-year (1970 - 2009) inventory using morphology (Marshall et al. 2009) revealed 2,423 taxa identified to a genus- or species-level among 30,000 specimens collected from Point Pelee and vicinity. After merging the two inventories, there were 3,217 genera/species combinations in the checklist with just 7.8% overlap (Table S2, doi:10.5883/DS-PPNP12). The overall taxonomic coverage includes 343 families, 597 subfamilies, 1,783 genera, 2,290 species and another 118 interim or uncertain species. While the study by Marshall et al. (2009) only examined insects, the present study examined four classes of arthropods. Only considering insects, the present inventory revealed more species of Trichoptera, Thysanoptera, and Psocodea. When all BINs are considered, the present inventory was biased toward Diptera and Hymenoptera where it collected 19.8% and 13.0% more respectively. By contrast, the diverse collecting methods employed by Marshall et al. (2009) yielded more Coleoptera, Hemiptera and Lepidoptera (64.6%, 31.6%, 22.2%). In total, the present effort added 780 taxonomic records to the checklist (Table S2; doi:10.5883/DS-PPNP12) which included 523 new species, 396 new genera, 91 new subfamilies, and 86 new families.

## Discussion

This paper describes the steps involved in moving from specimen collection through DNA barcode analysis to a summary of species, their abundances and associated diversity metrics. Aside from enabling a rapid, inexpensive assessment of terrestrial arthropod diversity, this approach aids extension of the DNA barcode reference library.

### Capturing Presence and Abundance

The current pipeline overcomes several barriers that usually constrain surveys of arthropod diversity. Most importantly, DNA barcoding minimizes the time demand on taxonomic experts by automating the identification of specimens that belong to species in the reference library (deWaard et al. 2009, Telfer et al. 2015). As a consequence, taxonomic advice is only required when a new BIN is encountered. The use of BINs also streamlines barcode workflows. For example, although specimen images are important to identify new BINs, a few representatives of each taxon are adequate. Similarly, a carefully edited bidirectional sequence is required for each new BIN, but a unidirectional sequence is perfectly adequate for BIN assignment since intraspecific variation within a population is low (Bergsten et al. 2012). For instance, two BINs of *Empoasca* (Hemiptera: Cicadellidae) were represented in the Point Pelee collection by 555 (BOLD:AAG2868) and one (BOLD:ACZ4093) specimens respectively. Just a few representatives of the abundant BIN were imaged and bio directionally sequenced, but every specimen could be identified by unidirectional analysis. Aside from allowing the strategic deployment of analytical effort, the key advantage of DNA barcoding lies in its capacity to allow technicians with no taxonomic training to generate the species abundance data needed for most diversity indices (Magurran 2004). As well, abundance data are valuable to employ functional traits to quantify ecosystem processes and services (e.g., Devictor et al. 2010). In addition, abundance data coupled with sequence information on each specimen allows genetic diversity to be quantified (Miraldo et al. 2016), which enables follow-up examinations such as probing the correlation between species richness and genetic diversity (Vellend 2005).

### Assembly of Resources

As evidenced by our study at Point Pelee, this approach generates a taxonomic inventory, an image library, a DNA archive, sequence data and specimens with associated collection data, information that can be shared through diverse online portals (e.g. Telfer et al. 2015). It also expands the DNA barcode reference library in a more cost-effective way than by analyzing legacy specimens because of their degraded DNA (Hebert et al. 2013; Prosser et al. 2016). As well, the analysis of newly collected specimens permits supplemental investigations, such as genome size determination (Hanner and Gregory 2007) and stable isotope analysis (Dittrich et al. 2017). The barcode library has utility beyond species identification, including the reconstruction of community phylogenies (e.g. Boyle & Adamowicz 2015) for studying the structure and assembly of biological communities, as well as for flagging new species (e.g. van Nieukerken et al. 2015) and new occurrence records (Fernandez-Triana et al. 2014).

### Protocol Refinements

The present method has gained wide adoption (Perez et al. 2015, http://www.globalmalaise.org; Zlotnick et al. 2015; Steinke et al. 2017), being employed in several studies (Bukowski et al. 2015; D’Souza et al. 2015; Kohn et al. 2015; Mazumdar et al. 2015; Aagaard et al. 2017; Geiger et al. 2016; Hebert et al. 2016; Wirta et al. 2016). In fact, 2.8 million specimens have now been processed using this method. This work has led to one important improvement — a standard primer cocktail, C_LepFolF and C_LepFolR (Folmer et al. 1994; Hebert et al. 2004) that can be used for all arthropods, simplifying consolidation and sequencing. The present protocol generates high quality barcode records for approximately $3 a specimen with about two thirds of the cost derived from Sanger sequencing. A substantial reduction in analytical costs can be achieved by shifting to a high-throughput sequencing (HTS) platform. The Illumina MiSeq and Ion S5 platforms reduce sequencing costs four-fold (e.g. Shokralla et al. 2014, 2015, Meier et al. 2016, Morinière et al. 2016) while the PacBio Sequel System reduces them 40-fold (Hebert et al. 2017).

### A Global Terrestrial Arthropod Monitoring Network?

The deployment of an extensive network of Malaise traps is relatively inexpensive, as evidenced by past deployments in national parks (Perez et al. 2015), schoolyards (Steinke et al. 2017), and backyards (Zlotnick et al. 2015). Once the present approach has been integrated with HTS, the mass samples resulting from a broad trap network will deliver accurate occurrence data while extending the barcode reference library. By monitoring biodiversity on a massive scale, this activity would advance each country’s capacity to deliver factually-based reports on the status of biodiversity as required to meet the Convention on Biological Diversity’s Aichi Targets of the Strategic Plan for Biodiversity 2011–2020 (https://www.cbd.int/sp/targets/).

## Author Contributions

J.R.D., S.L.D., J.E.S., E.V.Z. and P.D.N.H conceived and designed the study. S.L.D., C.N.S, R.M., J.T.A.M., A.D.Y. performed the experiments. M.R.Y., V.L.B., J.E.S. and K.P. analyzed the data. N.V.I., S.R., S.N., C.S. and A.C.T. contributed reagents/materials/analysis tools. J.R.D., S.L.D., J.E.S., V.L.B. and P.D.N.H. wrote the paper. All authors read and approved the final manuscript.

## Acknowledgements

The Ontario Ministry of Research and Innovation enabled this study through grants in support of the International Barcode of Life project while the Canada Foundation for Innovation provided essential infrastructure to the Centre for Biodiversity Genomics (CBG, http://www.biodiversitygenomics.net). We particularly thank Anne McCain Evans and Chris Evans for generously supporting our research program. Our work depended heavily on analytical support provided by the Barcode of Life Data Systems (BOLD, http://www.boldsystems.org). We thank John Waithaka, Heidi Brown, Tammy Dobbie and other Parks Canada staff who facilitated both permit acquisition and specimen collections. We also thank colleagues at the CBG including S. Bateson, G. Blagoev, A. Borisenko, V. Campbell, C. Christopoulos, J. Gleason, K. Hough, L. Lu, R. Manjunath, M. Milton, S. Pedersen, S. Prosser, J. Robertson, D. Roes, D. Steinke, A. Stoneham, J. Topan, C. Warne, and C. Wei.

## Data accessibility

Data analyzed in this paper are available in several repositories and formats: all specimen and sequence data are available on BOLD in the dataset DS-PPNP12 (dx.doi.org/10.5883/DS-PPNP12); an abridged version of the data is available in Appendix S1, which includes the GenBank accessions for the records on NCBI; the occurrence data are included in a Darwin Core Archive available on GBIF through the Canadensys portal at doi:10.5886/qzxxd2pa; and a BOLD checklist of the updated Point Pelee National Park arthropod inventory is accessible at doi:10.5883/DS-PPNP12.

## Supporting Information

**Fig. S1** Neighbor-Joining tree based on sequence divergences at COI (K2P distance model) for one representative of all 2,254 BINs.

**Fig. S2** Image library matching the COI Neighbor-Joining tree of BIN representatives. In a few instances, an image for the BIN representative was unavailable because the specimen was not recovered after DNA extraction. In these cases, an image of a different representative of the same BIN from another site was chosen, or in rare cases, from the nearest neighbor BIN (as marked below the image).

**Appendix S1** BOLD and GenBank accessions, as well as BIN assignments and collection details for the 19,185 arthropods from Point Pelee National Park.

**Appendix S2** Comparison of genera and species found at Point Pelee National Park by morphological (Marshall et al. 2009) and DNA barcode inventories.

